# Anterior forebrain pathway in parrots is necessary for producing learned vocalizations with individual signatures

**DOI:** 10.1101/2023.05.04.539392

**Authors:** Zhilei Zhao, Han Kheng Teoh, Julie Carpenter, Frieda Nemon, Brian Kardon, Itai Cohen, Jesse H. Goldberg

## Abstract

Parrots have enormous vocal imitation capacities and produce individually unique vocal signatures. Like songbirds, parrots have a nucleated neural song system with distinct anterior (AFP) and posterior forebrain pathways (PFP). To test if song systems of parrots and songbirds, which diverged over 50 million years ago, have a similar functional organization, we first established a neuroscience-compatible call-and-response behavioral paradigm to elicit learned contact calls in budgerigars (*Melopsittacus undulatus*). Using variational autoencoder-based machine learning methods, we show that contact calls within affiliated groups converge but that individuals maintain unique acoustic features, or vocal signatures, even after call convergence. Next, we transiently inactivated the outputs of AFP to test if learned vocalizations can be produced by the PFP alone. As in songbirds, AFP inactivation had an immediate effect on vocalizations, consistent with a premotor role. But in contrast to songbirds, where the isolated PFP is sufficient to produce stereotyped and acoustically normal vocalizations, isolation of the budgerigar PFP caused a degradation of call acoustic structure, stereotypy, and individual uniqueness. Thus the contribution of AFP and the capacity of isolated PFP to produce learned vocalizations have diverged substantially between songbirds and parrots, likely driven by their distinct behavioral ecology and neural connectivity.

## Introduction

The ability to learn vocalizations and use them for social communication was a key milestone in the evolution of human language ^1^. This trait, rare among non-human animals, is observed in three clades of birds: oscine songbirds, parrots, and hummingbirds ^2–4^. Interestingly, all of these clades evolved nucleated forebrain circuits, or ‘song systems’. Superficially, these song systems appear similar because they share a posterior cortico-cortical pathway and an anterior forebrain pathway comprised of a basal ganglia (BG) thalamocortical loop ^5,6^, but because these clades diverged over 50 million years ago and have non-vocal learners in their lineages, it remains unknown if the functional organization of these pathways are conserved ^1,4^.

The song system has been extensively studied in three songbird species: zebra finches, Bengalese finches, and canaries ^7–10^. Conceptually similar results across these species support a conserved logic where anatomy maps to function ^5,6^ (Figure 1A). The output of the song system, RA (robust nucleus of the arcopallium), projects to brainstem motor neurons and receives two major inputs: 1) from HVC (used as proper name), the output of the posterior forebrain pathway (PFP); and 2) from LMAN (lateral magnocellular nucleus of the anterior nidopallium), one of the two output nuclei of the anterior forebrain pathway (AFP) that forms a BG-thalamocortical loop. In songbirds, the AFP and PFP have very distinct roles in adult song production. Lesions or inactivations of HVC degrade the acoustic structure of adult song ^9,11^, revealing that the AFP in isolation produces abnormally variable vocalizations even in adults. In contrast, AFP inactivations result in songs with preserved acoustic structure but increased stereotypy ^12–17^, revealing that the PFP in isolation is sufficient for the production of learned and individually unique vocalizations.

**Figure 1.**
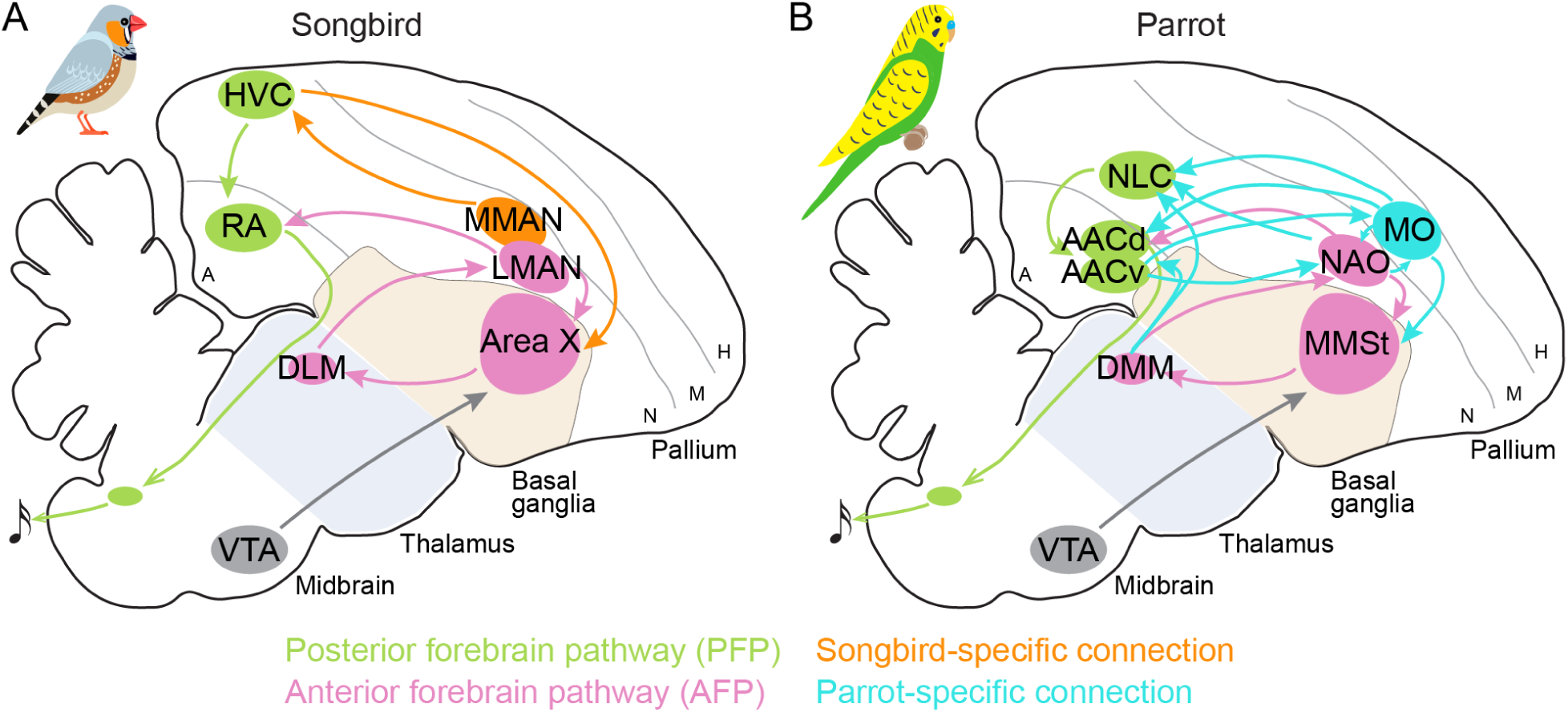
Vocal learning circuits in songbirds and parrots. (A) Sagittal view showing the posterior forebrain pathway (green) and anterior forebrain pathway (pink) in the zebra finch. Orange circles and lines indicate the songbird-specific vocal nuclei and connections. Abbreviations: H, hyperpallium; M, mesopallium; N, nidopallium; A, arcopallium; RA, robust nucleus of the arcopallium; HVC, proper name; LMAN, lateral magnocellular nucleus of the anterior nidopallium; MMAN, medial magnocellular nucleus of the anterior nidopallium; DLM, medial portion of the dorsolateral thalamus; VTA, ventral tegmental area. (B) Vocal learning circuits in budgerigars also have the posterior (green) and anterior (pink) pathways. Cyan circles and lines indicate the parrot-specific vocal nuclei and connections. The shell structures that surround the vocal nuclei are not depicted. Abbreviations: AAC, central nucleus of the anterior arcopallium; NLC, central nucleus of the lateral nidopallium; MO, oval nucleus of the anterior mesopallium; NAO, oval nucleus of the anterior nidopallium; MMSt, magnocellular nucleus of the medial striatum; DMM, magnocellular nucleus of the dorsal thalamus.

To test if these pathway-specific functions generalize across distantly related vocal learners, we studied adult vocal production in the budgerigar (*Melopsittacus undulatus*), a member of the parrot family. Anatomical studies show complex similarities and differences between the parrot and songbird song systems (Figure 1) ^18–27^. First, AAC (central nucleus of the anterior arcopallium) may be analogous to RA because it is similarly positioned in the posterior forebrain, projects to motor neurons in the brainstem, and receives inputs from both a posterior and frontal pathway ^18–22^. Second, NLC (central nucleus of the lateral nidopallium) has been proposed to be analogous to HVC because of its posterior location, its prominent projection to AAC, and its inputs from high-order auditory areas ^22,24,25^. Third, MO (oval nucleus of the mesopallium) and NAO (oval nucleus of the anterior nidopallium) share anatomical features with LMAN, because they are similarly positioned in the anterior forebrain and project to both AAC (output of song system) and BG, forming a BG-thalamocortical loop similar to the songbird AFP.

Yet closer examination of the song systems shows key differences that may have major functional importance (Figure 1) ^18–27^. First, the budgerigar song system has substantially more connectivity. For example, the songbird AFP has two output nuclei, LMAN and MMAN (medial magnocellular nucleus of the anterior nidopallium). MMAN and LMAN project to HVC and RA, respectively, and have limited internuclear connectivity ^28^. But parrot AAC receives inputs from two frontal structures (MO and NAO) that immediately neighbor and heavily interconnect with each other ^20,22,23,25,29^. The parrot song system also has substantially more long-range recurrence (Figure 1). For example, while LMAN and MMAN project to RA and HVC, respectively, MO and NAO both project to both AAC and NLC, forming a feedforward loop ^20,22,25,26^. Further, while RA projects primarily to the brainstem and thalamus ^30^, with minimal projections back to HVC and, as far as is known, none back to LMAN, AAC receives a direct thalamic projection and has substantial recurrent connectivity with its forebrain inputs ^19,20,22,26^. Finally, each ‘core’ nucleus of the parrot forebrain nuclei is surrounded by ‘shell’ tissues that differ in connectivity and molecular markers ^26^. In songbirds, shell tissue exists around some nuclei, but are not substantially interconnected with the cores in adults ^31^. Yet in parrots, shell tissue of MO in the forebrain pathway and AAC in the posterior pathway forms extensive local recurrent connections with the cores as well as long-range recurrent connections to other nuclei ^26^.

The functional consequences of these anatomical similarities and differences remain unclear. Previous functional studies in budgerigars focused primarily on the posterior NLC-AAC pathway. Bilateral lesions of AAC disrupted the acoustic structure of learned vocalizations ^21,32^, while partial lesion of NLC affected amplitude modulation ^33^, consistent with the idea that PFP is necessary for vocal production in parrots, just as in songbirds. However, it remains unknown whether parrot PFP is sufficient for the production of learned vocalizations as in adult songbirds, or AFP also plays a necessary role. To our knowledge, only one study conducted lesions of MO in one bird and found that frequency modulation of contact calls was initially normal but gradually lost over several days ^21^. The authors proposed that MO may be important for learning and maintenance of contact calls but not in vocal production. Given the low sample size of this study, the limited histological data, and the limited analyses of a small vocalization dataset, as well as the possibility that the gradual degradation of call acoustic structure was a by-product of their aspiration lesion spreading over time, we sought to revisit the functions of AFP and PFP in parrot vocal production.

We examined the functions of parrot AFP and PFP in adult vocal production at the pathway level, and compared the results to those in songbirds. In adult songbirds, lesions or inactivations of AFP achieve two goals: they clarify what aspects of vocalization are lost with AFP eliminated and also what aspects of vocalizations can be driven by the PFP alone. In zebra finches and Bengalese finches, LMAN inactivations in adult birds cause an immediate reduction in vocal variability but leave the mean acoustic structure largely intact ^12–17^, showing that PFP is sufficient to drive individually unique songs in these songbird species. Here we adopt the same logic and method to adult budgerigars. We test if parrot PFP, isolated by acute AFP inactivations, is sufficient to produce intact vocalizations that are already learned. If AFP-inactivated budgerigars can still produce stereotyped and individually unique learned vocalizations, as has been observed in songbirds, then key pathway-specific functions are shared between songbirds and parrots despite the anatomical differences. Alternatively, if AFP inactivations cause a degradation of learned vocalizations and loss of individual uniqueness, then this would indicate that the distinct evolutionary histories, ecologies, and known anatomical differences have resulted in substantial divergence of pathway-level functions.

## Results

### A behavioral paradigm to elicit thousands of learned contact calls per bird

Studying the detailed acoustic structure of vocalizations and how they may change with neural manipulations requires analysis of large datasets, ideally thousands of calls per bird ^34–36^. We set out to establish a behavioral paradigm that would provide such a dataset even in single-housed birds with pristine acoustic isolation. We focused on contact calls, a major type of learned vocalizations in budgerigars, that are used to coordinate flock behavior during flight and to localize individuals in a colony ^37^. To sample contact calls efficiently, we developed a call-and-response behavioral paradigm. We first recorded colony noise from a social group (5 males, 3 females) and divided the recording into two-minute segments. We then isolated individuals in a sound-proof box and used directional microphones to record their vocalizations (Figure 2A). In a given trial, one segment of colony noise was randomly selected and played to the bird through a speaker, followed by a silent inter-trial interval (average 10 mins). Isolated birds responded to colony noise with the production of contact calls so reliably that we were able to obtain hundreds to thousands of contact calls per day with this method (counts/day range: 180-1612, n=8 birds, Figure 2B).

**Figure 2.**
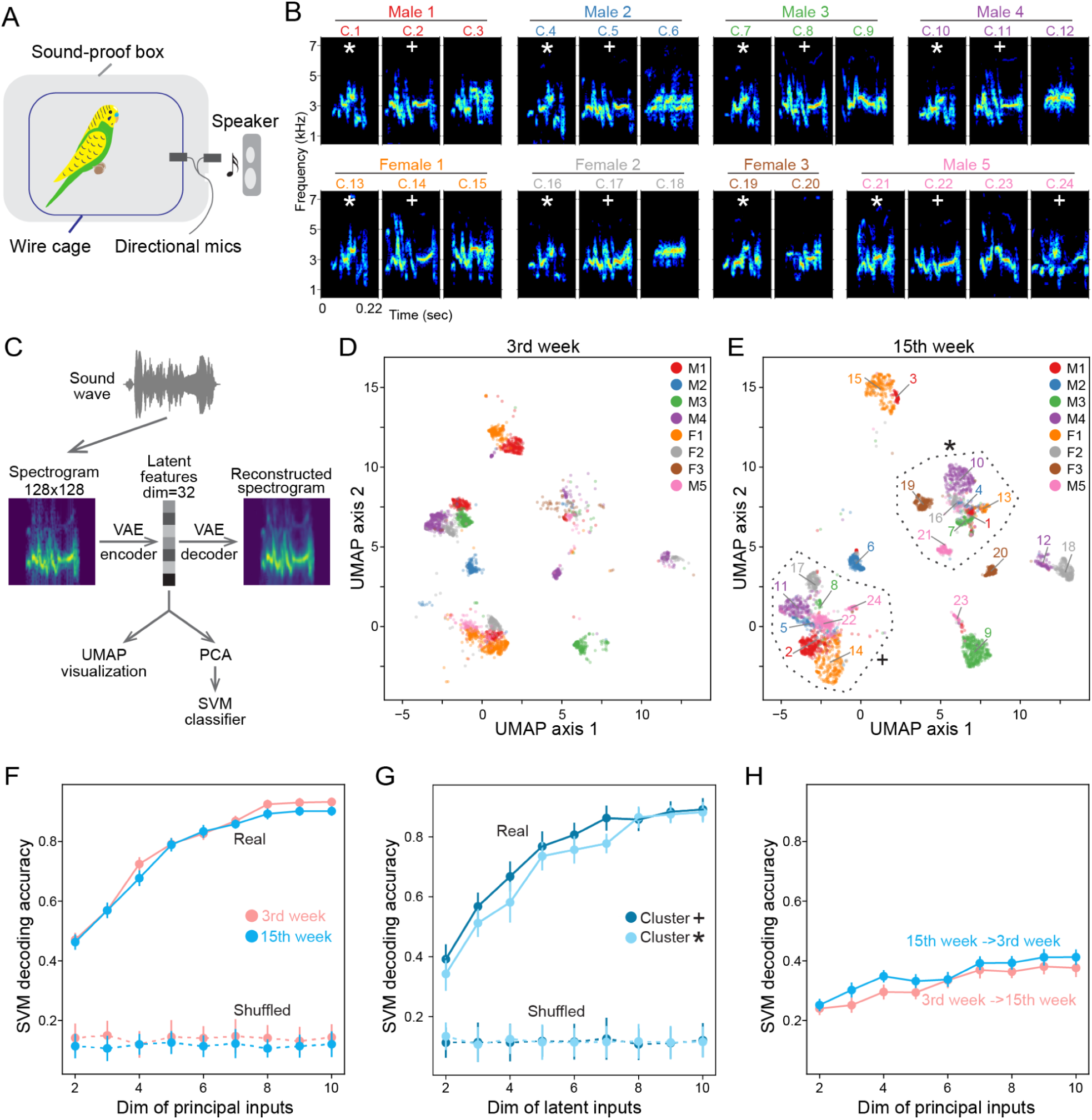
Budgerigar contact calls contain individual signature. (A) Schematics showing the recording setup. Two directional microphones point in the opposite direction to capture sound from inside or outside the box. (B) Spectrograms of example contact calls from a group of 8 budgerigars (5 males, 3 females) at week 15 since assimilation. All plots have the same scale on the x- and y- axis. Animal and cluster labels above each spectrogram correspond to labels in panel *E*. Asterisk symbol marks a call type that was shared across all 8 individuals, while plus symbol marks one that was shared across 7 individuals. (C) Pipeline for analyzing contact calls (D-E) UMAP embedding of contact calls at week 3 (D) and week 15 (E) since group assimilation. Two datasets were trained through the same VAE and embedded in the same UMAP space, but shown separately for clarity. Each dot is a call, colored by caller identity. A maximum of 600 calls were shown for each individual. Animal and cluster labels in (E) correspond to spectrograms in panel *B*. Dashed lines in (E) circled the two major clusters that were shared across individuals and marked by an asterisk or plus symbol in panel *B*. (F) Accuracy of SVM classifiers that decode individual identity from call latent features as a function of input dimensions. Solid lines are models trained on real data, while dashed lines are models trained on shuffled data. (G) Accuracy of SVM classifiers that were trained and tested solely on the calls that belong to the two large shared clusters (marked by asterisk and plus symbol in panel B and E). (H) Accuracy of SVM classifiers that were trained on one time point and used to decode individual identity at a different time point.

### A variational autoencoder identifies individual signatures even within shared call types

With this large dataset, we next developed machine learning methods to quantify the acoustic structure and individual uniqueness of budgerigar contact calls. Previous studies have shown that contact calls are learned and continuously adjusted even in adulthood ^38–42^. A well-characterized phenomenon is the convergence of contact calls within a social group ^39–41^. While call convergence may be useful to signal group affiliation, it also poses a challenge for social communication—how can group members signal individual identity if their contact calls become highly similar to each other? It remains unclear if and how individuality persists in the converged contact calls ^38,39,42^.

We wondered if even seemingly similar calls between affiliated birds exhibit enough acoustic distinctiveness as a prerequisite for individual discrimination. Examining this possibility requires sensitive methods to compare contact calls between individuals, beyond the traditional methods that were based on visual inspection or descriptive acoustic features ^39–42^. We therefore adopted a recently developed method ^36^, variational auto-encoder (VAE), to analyze contact calls in an unsupervised manner (Figure 2C, Methods). Briefly, vocalizations were first converted to spectrograms, then an encoder-decoder network was trained to reconstruct the original spectrograms from low-dimensional latent features. These latent features were used for downstream visualization with UMAP (uniform manifold approximation and projection) and for quantitative classification of uniqueness with SVM (support vector machine).

Using these methods, we set out to quantify the level of individuality in contact calls after convergence. First, we confirmed that individuals in our recorded group produced both unique calls as well as shared calls that exhibited highly similar acoustic structures ^39–41^, as shown by the spectrograms (Figure 2B) and clustering in the UMAP space (Figure 2D-E). Second, closer examination of the UMAP clustering pattern suggested that calls from individuals occupied distinct parts of the large clusters of converged calls, consistent with the existence of individuality even after call convergence. We quantified the level of individuality by training SVM classifiers to decode individual identity from call latent features ^43^. Unsurprisingly, as we increased the dimension of principal inputs, i.e. used more information from the spectrogram, the decoding accuracy gradually improved (Figure 2F). With eight or more principal inputs, the SVM correctly predicted bird identity more than 90% of the time. This accuracy was significantly higher than a previous study that used summary acoustic metrics ^42^ (∼40%), confirming that spectrograms contained extra information for discrimination not captured by acoustic feature vectors ^36^. Lastly, we restricted the analysis to the two major call types (Figure 2B, 2E, marked by asterisk and plus symbols) that were shared across most individuals, and found that the SVM still decoded bird identity ∼90% of the time (Figure 2G). Together, these results demonstrate that a high level of discriminatory information persists in budgerigar contact calls even for the shared call types and set the stage for future behavioral experiments to test if the information is used to allow for individual recognition in a social group.

### Group vocalizations dynamically change over time

A proposed function of contact calls is to signal affiliation among birds, which, in the wild, can change over time ^37,44–46^. Call matching would be a more honest signal of affiliation if calls dynamically changed over time, requiring ongoing social interactions and imitation ^40,44^. To test if our analyses could capture this dynamism of budgerigar contact calls, we compared contact calls of each individual in the group produced at week 3 (Figure 2D) and week 15 (Figure 2E) following the group assimilation. We trained a single VAE on the contact calls produced at these two time points and visualized them using UMAP (Figure 2D-E). We found that the positions of contact calls in the UMAP space were not static but exhibited drift over time, consistent with dynamic changes of call acoustic structure ^39^. We then quantified time-dependent changes in call structure by examining how well an SVM classifier trained on calls at week 3 could decode calls at week 15, and vice versa (Methods). Decoding accuracy dropped from ∼90% to ∼40% (Figure 2H), suggesting that the acoustic structures of contact calls changed significantly over the 12 weeks. These confirmations of past findings with new analytical methods applied to large call datasets lays the foundation for studying how brain inactivations affect call acoustic structure.

### AFP-inactivated budgerigars maintain the ability to respond to stimulus calls

We next wondered how calls are produced by the budgerigar song system. Reverse microdialysis is an established method in songbirds that can transiently and reversibly inactivate brain regions over hour timescales ^10,30,47^. We implanted custom microdialysis probes bilaterally at the center of the MO/NAO complex to inactivate both nuclei at once (Figure 3A-B, Methods), which is necessary for studying the capacity of isolated PFP. After birds recovered from surgery, we infused saline (PBS) or tetrodotoxin (TTX 50 µM) into the probe on alternating days to compare their effects on contact calls in the call-and-response paradigm. Importantly, we confirmed with electrophysiological experiments in anesthetized animals that the diffusion of 50 µM TTX caused fast inactivation in brain regions ∼1 mm from the probe location, large enough to cover the MO/NAO complex. We also confirmed that activity in distant structures over four millimeters away, such as the NLC-AAC pathway, remained intact (Figure S1). These calibration results are consistent with our past calibrations using these identical probes ^11,30^.

**Figure 3.**
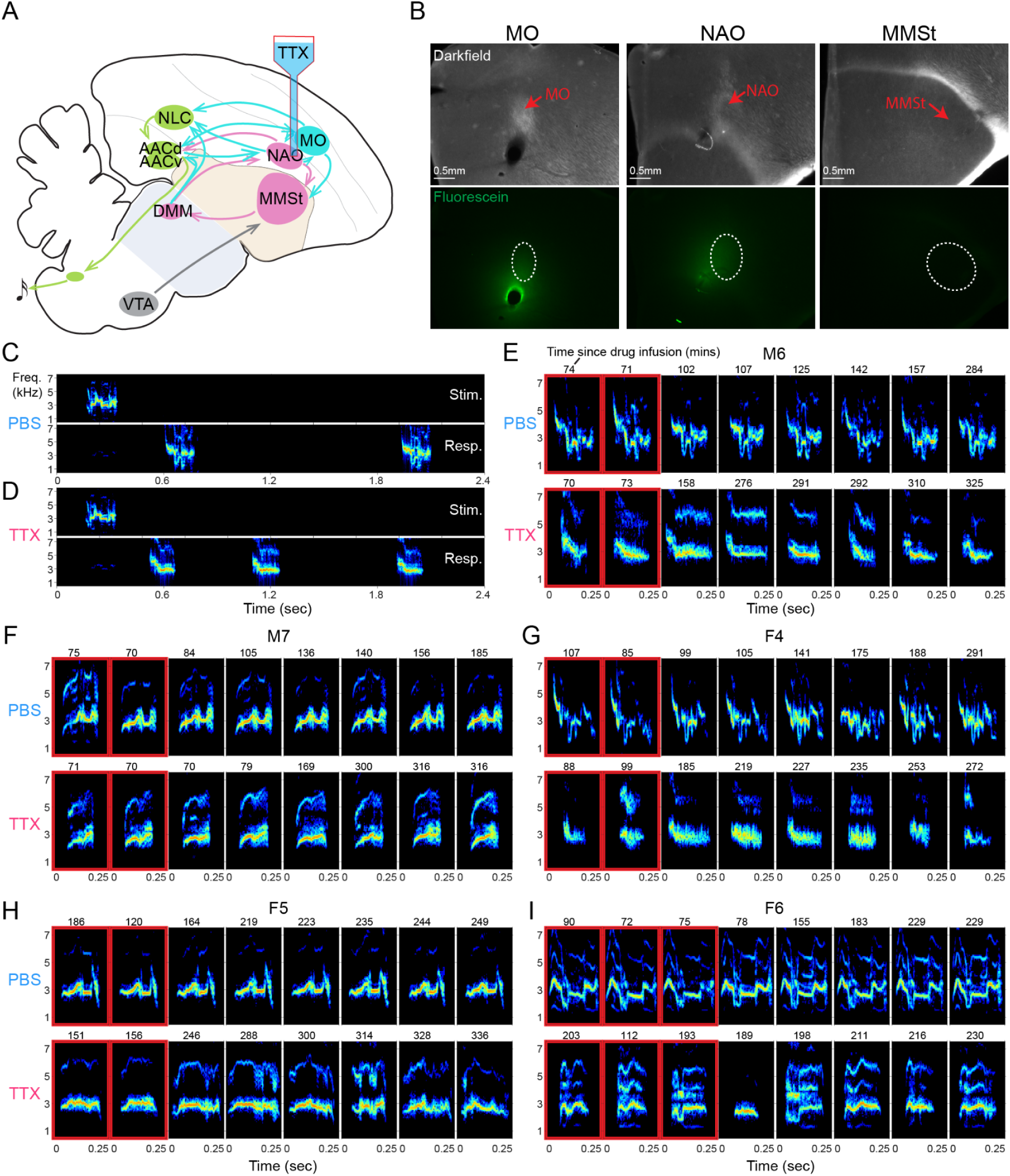
Frontal cortical inactivation abolishes pitch modulation in contact calls. (A) Schematics of the inactivation experiments. Reverse microdialysis probes were implanted in the center of the MO/NAO complex and infused with either 50 micromolar TTX or PBS. (B) Histological confirmation of implant site in one representative bird. At the end of experiments fluorescein was infused into the probes to visualize diffusion and probe placement. Fluorescein covered both MO and NAO. Dashed circles at the bottom row indicate the boundaries of corresponding nuclei. (C-D) Example trials from one bird under the PBS (C) and TTX (D) condition. Top rows show spectrograms of the stimuli, bottom rows show time-aligned responses. Y-axis is frequency in kHz. (E-I) Example spectrograms from 5 birds under the PBS (top row) and TTX (bottom row) condition. The first 2-3 spectrograms at each row in the red boxes are the first calls produced during each PBS or TTX infusion replicate. Other spectrograms were selected randomly from all infusion replicates and sorted in chronological order. M6-M7, males; F4-F6, females. Y-axis is frequency in kHz. Numbers above each spectrogram indicate minutes since drug infusion. Note that because the first calls given by a bird at different infusion replicates might vary in time, the randomly selected calls might appear earlier than the first calls in some infusion replicates.

To sample contact calls, we used the same call-and-respond behavioral paradigm (Figure 2A), but played individual calls instead of colony noise to better examine the temporal relationship between stimulus and response (Figure 3C-D). To test the capacity of the intact and isolated PFP in producing calls with clearly distinct acoustic structures, we chose five adult budgerigars unknown to one another with clearly distinguishable contact calls (Figure 3E-I). Unlike in songbird models where vocal learning is largely sexually dimorphic, both male and female budgerigars readily learn and produce contact calls and possess similar nucleated song systems ^22,37,41,48^. Because we did not observe obvious sexual differences in the effects of AFP inactivation, the following results combine data from males and females.

With MO/NAO inactivated, all birds retained their ability to respond to call stimuli (Figure 3C-I). Response rate significantly decreased in three of five birds (Figure 4A). Response latency changes were variable: two birds responded slower, one bird faster, while the remaining two birds exhibited no change in latency (Figure 4B). Together these data show that the AFP-inactivated budgerigars maintained the ability to respond to stimulus calls through the activity of PFP alone.

**Figure 4.**
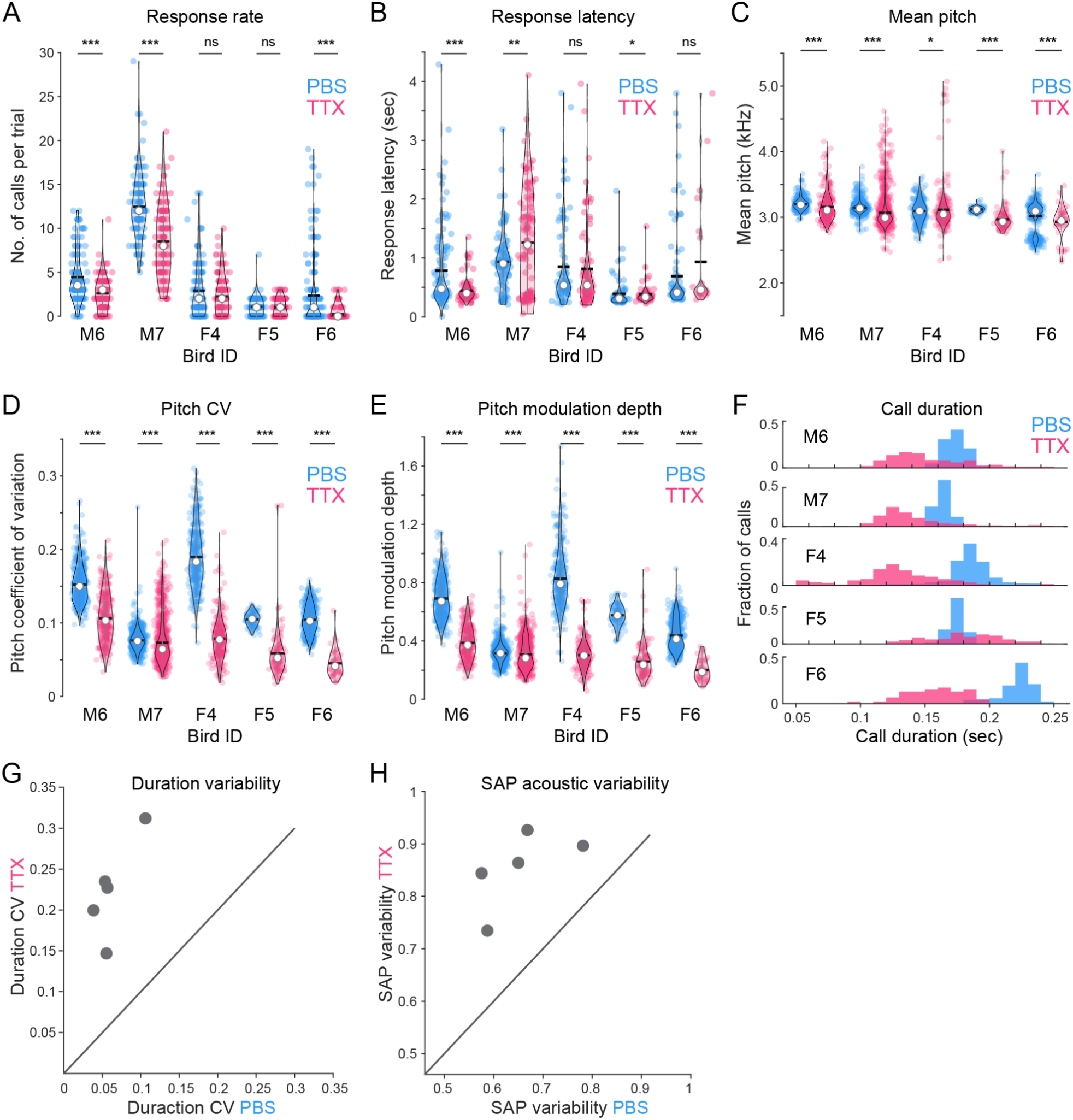
Frontal cortical inactivation increases variability of contact calls. (A-E) Compare behavioral trial statistics (A-B) and acoustic features (C-E) between the PBS (blue) and TTX (red) condition. Each colored dot is one trial (A-B) or one call (C-E). The violins indicate the shape of distribution. The black line and white dot indicate the mean and median. P-value: *<0.05, **<0.01, ***<0.001, ns not significant, with Mann-Whitney U tests. (F) Histograms of contact call duration under the PBS (blue) and TTX (red) condition. (G) Variability of call duration quantified by coefficient of variation (CV). Each dot indicates data from one bird.(H) Variability of acoustic features calculated by Sound Analysis Pro (SAP). Each dot indicates the averaged dissimilarity of 50 randomly selected pairs of calls in a bird under the PBS or TTX condition.

### AFP inactivation disrupts call acoustic structure and increases cross-rendition variability

We next examined the acoustic structure of contact calls before, during, and after AFP inactivations. AFP inactivations resulted in a subtle but significant decrease in the mean pitch of calls, but they remained in the normal species-typical frequency range of 2-4 kHz ^37^ (Figure 4C), suggesting that all birds were able to produce contact-call-like vocalizations. However, two aspects of call acoustic structure were dramatically disrupted. First, calls produced under the TTX condition exhibited significantly less pitch modulation compared to vehicle, and virtually all of the rapid pitch modulations were eliminated (Figure 3E-I). We quantified the level of frequency modulation within a single contact call by first estimating its pitch contour over time, then calculating the coefficient of variation and modulation depth [(max-min)/mean] of the pitch values (Methods). We found that both measures were significantly lower for the AFP-inactivated calls (Figure 4D-E, PBS:TTX ratio, 1.80±0.58 for coefficient of variation, 1.99±0.64 for modulation depth, p<0.001 for all five birds, Mann-Whitney U tests). Second, call durations were affected. Normal contact calls, and those observed with vehicle in the probes, exhibited durations between 0.15 to 0.25 sec with a narrow distribution, consistent with past studies (Figure 4F) ^37,39^. AFP inactivation resulted in a significant increase in call duration variability, including the production of abnormally short or long calls never seen in control conditions (Figure 3E-I, Figure 4F, e.g. long calls observed in bird M6 and F5).

In zebra finches and Bengalese finches, AFP inactivation results in the preservation of learned syllable acoustic structure and an increase in syllable stereotypy across renditions ^12–17^. To test if inactivations of parrot AFP have a similar effect, we quantified cross-rendition variability, the variability in acoustic structures of the same syllable across different renditions, of contact calls using the same metrics previously used in songbirds ^13,17^. First, the variability of call durations, as measured by coefficients of variation, was significantly larger for the AFP-inactivated calls over the entire experimental period (Figure 4G, p=0.0011, paired t-test; Figure S2). Second, the variability in phonology, as quantified by the syllable level feature vectors defined and calculated in Sound Analysis Pro software, was also significantly larger under AFP inactivation (Figure 4H, p=0.0027, paired t-test, Methods). Thus, in contrast to songbirds, AFP inactivation in adult budgerigars results in the degradation of call acoustic structure and a dramatic increase in cross-rendition variability.

To check if these defects were caused by impairments in vocal production or feedback learning as proposed by a previous MO lesion study where there was a day timescale delay between lesion and behavioral deficit ^21^, we next examined the timescale over which behavioral deficits were expressed relative to the infusion of TTX. We observed defects in pitch and duration in the first calls produced immediately after TTX infusion (Figure 3E-I, red boxes). The acoustic structure of contact calls gradually recovered over hours after TTX washout (Figure S3). These results support a direct role of parrot AFP in the production of learned vocalizations.

Lastly, we performed two control experiments to confirm the specificity of our AFP inactivations. First, because TTX may also affect the fibers of passage coursing near MO/NAO, in one bird we had the opportunity to infuse muscimol that achieves inactivation by recruiting local inhibitory neurons. We observed identical effects on call acoustic structure in this bird (Figure S4), but future studies with a larger sample size of muscimol inactivation are needed to rule out the possible effects caused by fiber of passage. Second, we inactivated PFP in three birds with the same reverse microdialysis method to test if the effects of AFP inactivations we observed could be simply explained by drug leaking into PFP (Figure S5). The acoustic structures of PFP-inactivated calls were also dramatically degraded, but in a way not identical to AFP inactivations, consistent with the electrophysiological calibration (Figure S1) and the idea that parrot AFP provides crucial and specific premotor inputs to PFP to drive the production of learned vocalizations.

### AFP inactivation reduces the individuality of contact calls

The significant loss of pitch modulation and increase of phonological variability with AFP inactivation may reduce individual signatures in contact calls. We first examined this possibility by embedding the normal and AFP-inactivated calls of all birds in the same UMAP space. Since normal calls of these stranger birds are visually distinct (Figure 3E-I), they should form well-separated clusters in the UMAP. Changes in acoustic structure induced by AFP inactivation will drive TTX calls to deviate from the normal clusters. However, the deviation could happen in two ways that lead to different outcomes on the individual signatures (Figure 5A). If AFP inactivation reduces individuality, calls from different birds should move away from their normal clusters and converge into one large centered cluster (Figure 5A, hypothesis 1). Alternatively, if AFP inactivation affected each bird in unique ways, individuality may be maintained, in which case the UMAP embeddings of the PFP-driven calls would still reveal clearly distinct clusters associated with different individuals (Figure 5A, hypothesis 2). Our data were consistent with hypothesis 1: contact calls of different individuals converged to a large centered cluster following AFP inactivation (Figure 5B, S6), suggesting that PFP-driven calls were more similar across individuals than the normal calls and that AFP activity contributes to individuality.

**Figure 5.**
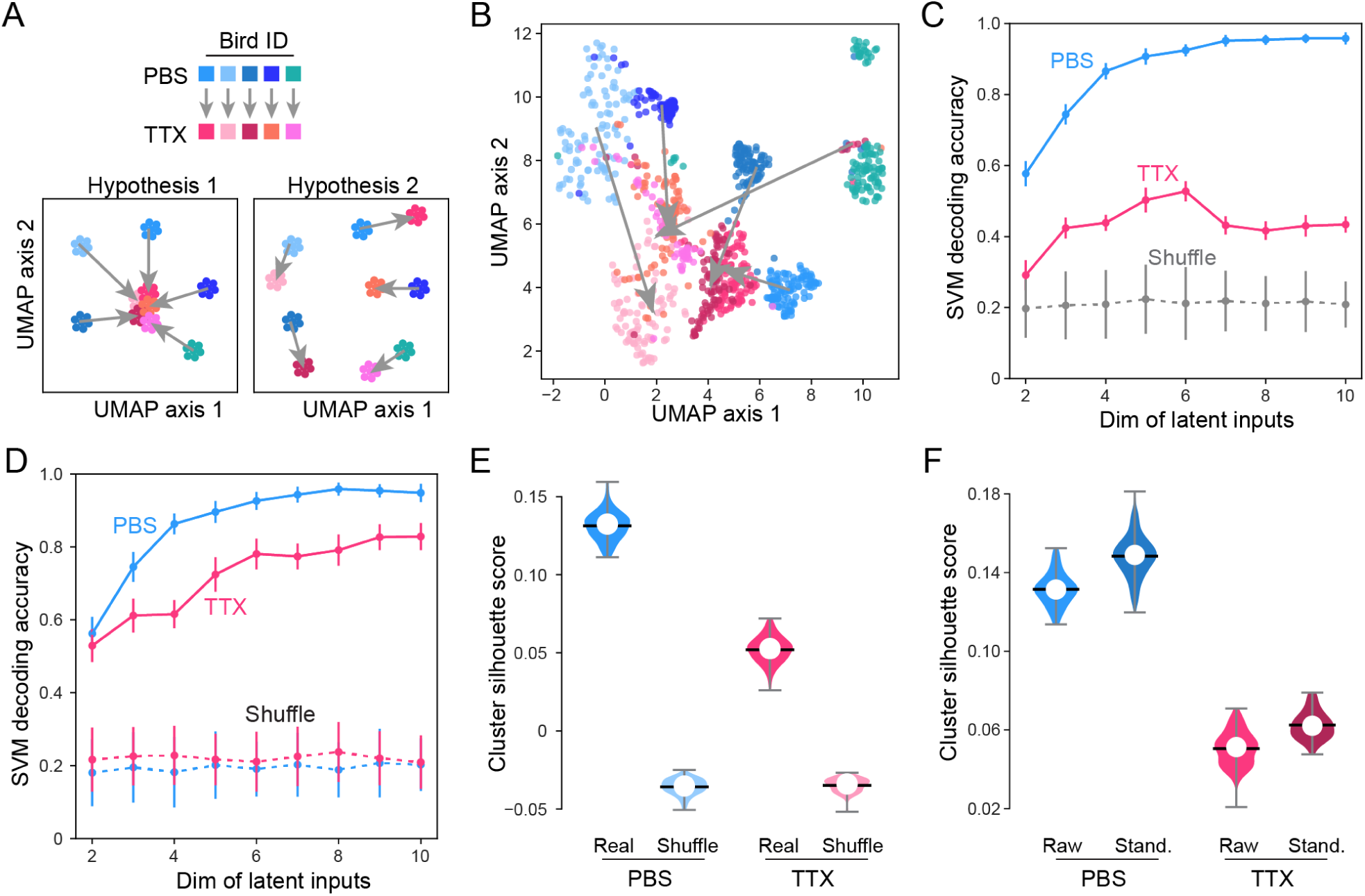
Frontal cortical inactivation reduces individual signatures in contact calls. (A) Hypotheses on how AFP inactivation would change the embedding of contact calls in the UMAP space. Clusters are colored according to individual identity and experimental condition. Gray arrows point from PBS to TTX condition in the same birds. Hypothesis 1: if AFP inactivation reduces individuality, calls from different birds should move away from their normal clusters and converge into one large centered cluster. Hypothesis 2: if AFP inactivation affected each bird in unique ways, individuality may be maintained, in which case the UMAP embeddings of the PFP-driven calls would still reveal clearly distinct clusters associated with different individuals. (B) UMAP embedding of contact calls. Each dot indicates a call, colored by caller identity and experiment condition. Gray arrows point from the centroid of PBS cluster to the centroid of TTX cluster for each bird. (C) SVM classifier accuracy as a function of input dimensions. The model was trained on the PBS condition, then used to decode individual identity at the TTX condition. Dots indicate the mean accuracy, vertical lines indicate the standard deviation. (D) Same as (C), but two models were trained separately for PBS and TTX. (E) Separability of contact calls produced by different individuals at the PBS and TTX condition, as quantified by Silhouette score (Methods). Equal number of calls were sampled from each individual for the analysis. The results of 100 sampling procedures are plotted. Colored violins indicate the shape of Silhouette score distribution. Black lines and white dots indicate the mean and median values, respectively. (F) Same as (E), but comparing the Silhouette scores between raw and standardized datasets where the cluster sizes are normalized for the PBS and TTX datasets.

Since UMAP is a non-linear method, large distances in UMAP space do not directly translate to large differences in acoustic space. To more rigorously quantify the loss of individuality with AFP inactivation, we first asked how much discriminatory information was lost for a given individual due to the degradation of acoustic structures with AFP inactivation. We trained a SVM classifier on normal calls, then used it to decode the caller identity in the TTX condition. The decoding accuracy dropped from ∼95% to ∼50% (Figure 5C), demonstrating a significant reduction, but not complete elimination, in individual identity information following AFP inactivation.

We next examined precisely how much individuality was intact in the group of PFP-driven calls. PFP-driven calls could still retain uniqueness due to individual differences in the biomechanics of the vocal organ and/or learned components retained in the NLC-AAC pathway. We trained two SVM classifiers, one for the vehicle and one for the TTX dataset, and found that the decoding accuracy on the TTX dataset was significantly lower (Figure 5D). We further quantified the separability of calls across individuals using a Silhouette score that contrasts intra- and inter-cluster distances, with larger scores meaning better separability (Methods). We calculated Silhouette scores in the PCA space of call latent features, where unlike UMAP the global distance measures are meaningful for comparison. We found that the Silhouette score was significantly lower for the TTX dataset (Figure 5E). The reduced discriminability under AFP inactivation could be caused by degradation of acoustic structures and/or increased cross-rendition variability. To quantify their contributions, we standardized the cluster sizes of each individual in the PCA space to control for the cross-rendition variability (Methods) and found that the Silhouette scores increased for both datasets (Figure 5F, PBS, 12.8%; TTX, 23.6%), indicating a contribution from the cross-rendition variability. After controlling for the increased variability, the discriminability was still lower for the TTX dataset, likely caused by the degraded acoustic structures. Together these data show that PFP-driven calls were less separable across individuals.

## Discussion

The diversity of vocal learning in different species of birds presents an ideal case to study the evolution of neural circuits underlying complex behaviors ^1–4^. Neural mechanisms of vocal learning have been extensively studied in several songbirds, but remain relatively unexplored in other clades. Here we tested songbird-inspired hypotheses in parrots regarding the distinct functions of the anterior and posterior forebrain pathways. First, we developed a call-and-response paradigm to generate thousands of calls per day per bird. Second, we used machine learning methods to visualize and analyze the individual uniqueness of learned calls, and discovered that even the converged calls exhibit robust individuality. Third, we used transient brain inactivation to test neural mechanisms of call production. As in songbirds, AFP inactivations caused an immediate change in vocal structure, including reduced within-syllable variability of pitch (Figure 4D-E), consistent with a premotor role of AFP in driving rapid pitch modulation of learned vocalizations ^13–17,49,50^. Yet in contrast to finches, where AFP inactivations preserve the acoustic structure and individual uniqueness but reduce the cross-rendition variability of learned songs, AFP inactivations in parrots caused a dramatic degradation of call structure, loss of vocal uniqueness, and increase of cross-rendition variability. These results show that the contribution of AFP and the capacity of the isolated PFP to produce learned vocalizations differ substantially across songbirds and parrots.

Our results complement previous lesion studies and provide novel insights into the functional organization of the parrot song system. First, effects of AFP inactivations differ clearly from those with AAC lesions ^21,32^. Although both manipulations disrupted frequency modulation of contact calls, fundamental frequency of AAC-lesioned calls dropped significantly to 2 kHz, deviating from the 3 kHz of normal and AFP-inactivated calls (Figure 4C). Call durations increased with AAC lesions, but durations of AFP-inactivated calls became more variable with mean values actually decreased in four out of five birds (Figure 4F). Second, effects of AFP inactivations also differ from those with NLC lesions and inactivations. Partial lesions of NLC left the frequency modulation intact but changed the amplitude modulation ^33^, while our AFP and PFP inactivations in the same birds showed distinct effects on acoustic structure (Figure S5). Third, a previous study which performed a frontal aspiration in one bird found that frequency modulation was normal immediately after surgery but gradually lost over several days ^21^. In contrast, our acute targeted AFP inactivations showed that the first calls produced immediately after TTX infusion already had defects in acoustic structures, consistent with a premotor role of the parrot AFP. We suspect that the gradual degradation of acoustic structures in the previous study may have been a by-product of their aspiration lesion spreading over time ^51^. Fourth, the effects of AFP inactivations are more consistent with defects in motor production, rather than action selection, because birds with vehicle in their probes never produced calls resembling those with AFP inactivations (Figure 3E-I). Together, our data suggest that unlike in adult songbirds, parrot AFP plays a direct and necessary role in the production of vocalizations that are already learned.

The distinct functions of AFP in parrots and songbirds such as zebra finches, Bengalese finches, and canaries likely arise from important differences at the behavioral and neural circuit levels. Behaviorally, parrots exhibit outsized vocal plasticity even as adults ^38–42^, and courtship success of an individual may depend on its capacity to imitate the continuously changing vocalizations of a potential mate ^52–54^. In contrast, in the most commonly studied songbird, the zebra finch, song may be evaluated on its stereotypy across renditions ^55^. Because female finches prefer songs that are more stereotyped, separating learning and production in different pathways may ensure that variability associated with practice and learning can be flexibly turned off in the courtship context ^56^. However, this clear learning-production separation may not be an ideal solution for open-ended learners like the budgerigars, because the birds need to constantly modify their vocalizations to match others in the group, as well as the evolving calls of a potential mate. It may be more beneficial to preserve a strong influence of the learning circuits on the motor output, so that vocalizations can be modified more rapidly. The distinct behavioral goals (produce stereotyped songs for courtship *vs* continuously modify vocalizations) may be one reason why AFP has different anatomy and different contributions to vocal production in adult finches and budgerigars. Importantly, future studies in songbirds with vocal complexity and learning capacity comparable to budgerigars such as European starlings ^57^ and superb lyrebirds ^58^ are needed to test whether the observed differences in AFP function are caused by evolutionary divergence between songbirds and parrots, or by the distinct features of vocal learning behaviors between budgerigars and finches. Furthermore, it remains possible that early in development certain circuit mechanisms are shared between finches and budgerigars, while distinct development trajectories drive the function of AFP to differ in adults. Future functional and neurophysiological studies that compare across developmental stages are necessary to examine this possibility.

At the neural level, though the songbird and parrot song systems are both nucleated and contain distinct anterior and posterior pathways, there are also important differences in the detailed connectivity likely related to the distinct effects of AFP inactivations ^18–27^. Foremost, while songbird and parrot AFPs are both BG-thalamocortical loops, they have distinct anatomical structures. Parrots have two frontal cortical nuclei, MO and NAO, that each project to both AAC and NLC ^18–26^. Songbirds also have two frontal nuclei, LMAN and MMAN, but they project exclusively to RA and HVC, respectively. Therefore an important caveat of our paper is that the AFP inactivations, though methodologically similar with respect to the microdialysis probes and TTX concentrations, may have been functionally distinct due to the distinct underlying anatomy. Specifically, our data do not rule out the possibility that either MO or NAO, or more specifically one of their projections to AAC, does in fact have LMAN-like functionality, including the production of vocal variability. Thus the more careful interpretation of our findings is not that the specific nuclei of the parrot and songbird AFP have distinct functions, but rather that the posterior pathways (HVC-RA and NLC-AAC) when isolated with AFP inactivations have distinct capacities for producing learned vocalizations. A second consideration is that MO/NAO inactivations will also inactivate their projections to NLC, raising the possibility that the loss of call acoustic structure was due to an impairment in the NLC function by losing the AFP inputs. Notably, AFP and NLC inactivations had different effects on call production and acoustics (Figure S5). Finally, each forebrain ‘nucleus’ of the parrot song system is recurrently interconnected with surrounding ‘shell’ tissue ^20,26^. Their roles in song learning and production are unclear, as the shell structures in songbirds appear to be activated not by singing behavior but by general movement, for example, in trained hopping tasks ^31,59^. In budgerigars, gestures such as headbobs are frequently coupled to the learned vocalizations as an integrated social display ^45^. It will be interesting for future study to examine the role of song nuclei and their surrounding shell structures in vocal-gestural coupling. More broadly, due to the more complex and extensive connectivity of the parrot song system, future electrophysiology studies that record from identified structures and pathways will clarify their distinct roles in social communication.

## Acknowledgements

We thank Michael Sheehan, Lindy McBride, and members of the Goldberg lab for comments. We thank Jack Bradbury and Sandra Vehrencamp for inspiration for the study.

## Author contributions

Z.Z. and J.G. designed the study. Z.Z. collected the data with help from F.N. and B.K.. Z.Z., J.C., F.N., and H.K.T. analyzed the data. H.K.T. received supervision from I.C.. Z.Z. and J.G. wrote the manuscript with help from all other authors.

## Inclusion and diversity statement

We support inclusive, diverse, and equitable conduct of research.

## STAR Methods

### RESOURCE AVAILABILITY

#### Lead contact

Further information and requests for resources and reagents should be directed to and will be answered by the Lead Contact, Jesse H. Goldberg.

#### Materials availability

This study did not generate any unique biological materials.

#### Data and code availability

Data generated in this study to reproduce the results can be accessed through Mendeley Data (10.17632/j8rpy4dc6c.1). Custom scripts can be obtained through the Lead Contact.

### EXPERIMENTAL MODEL AND STUDY PARTICIPANT DETAILS

Animal care and experiments were carried out in accordance with NIH guidelines and were approved by the Cornell Institutional Animal Care and Use Committee. Subjects used in this study were 10 male and 6 female budgerigars (*Melopsittacus undulatus*) that aged between 1 to 3 years. Birds were bred in our aviary or obtained from outside breeders (Magnolia Bird Farm, California; PetSmart, New York; OK Birds, Oklahoma). We used 8 birds for contact call quantification, 3 for determining anatomical coordinates, and 5 for chemical inactivation. Birds were group-housed in flight cages or individually housed in wire cages with a 12-12 light cycle. Food and water were provided *ad libitum*.

### METHOD DETAILS

#### Contact call playback

We assembled a group of 8 budgerigars (5 males and 3 females) from three separate cages. The assembled cage was placed in the aviary, roughly 5 meters away from other budgerigar cages, but not visually or acoustically isolated from other birds. We conducted two types of playback experiments. For the first type (Figure 2), we recorded the call repertoire of the 8 birds by playing noise of their own colony on-and-off in a trial structure. Colony noise was first recorded with a portable recorder (V618, EVIDA) inside their flight cage for 2-3 days. We removed the silent portion of the recording and chopped it into 2-min segments. We then isolated individual birds to a wire cage inside a sound attenuating box (Figure 2A). A flat-spectrum speaker (iLoud Micro Monitor, IK Multimedia) was placed outside the box (∼10 cm) and pointed toward an air-exchange hole on the box. We adjusted the speaker volume so the playback sound measured 53-55 dB inside the box. For each trial, we randomly chose a colony noise segment and played it to the bird through the speaker. The inter-trial interval followed a truncated exponential distribution (mean: 10 mins, range: 7.5-12 mins). Playback and response signals were recorded with directional microphones (Pro 35, Audio-Technica), bandpass-filtered (30-15,000 Hz) and amplified with a preamplifier (XL48, Midas), and digitized through a DAQ (USB6210, National Instruments) before sending to the computer. We wrote custom LabView codes to control the playback and recording. Each bird was recorded for 2 days.

For the second type of playback (Figure 3-5), we recorded responses to individual contact calls with the same hardware setup, but from 5 stranger birds that were unfamiliar with the stimulus calls. Because our inactivation experiments focused on the production of contact calls, we tried to minimize the learning component by isolating the birds from the group and individually housing them in sound-proof boxes, without direct interaction with other birds during the entire experimental period. We first constructed a pool of stimulus calls from the colony-noise responses of the 8 birds in the behavioral experiments (Figure 2) and normalized the amplitude of all calls to the same value. For each trial, we randomly selected one caller and one call from its repertoire, then played it three times to the experimental stranger bird with intervals uniformly sampled between 2-5 sec. The inter-trial interval followed a truncated exponential distribution (mean: 5 mins, range: 4-6 mins).

#### Surgery and histology

To perform surgery and histology in the budgerigar, we adopted procedures that are commonly used in the zebra finch. All surgeries were performed under isoflurane anesthesia. Origin for the anterior/posterior and medial/lateral axis was established relative to the sagittal sinus. Brain surface after removing dura mater was used to zero the dorsal/ventral axis. We adjusted the head angle of the bird until the anterior skull was horizontal, which was referred to as the zero-degree angle. To access a brain region, we removed the first layer of skull with a cranial drill, the second layer of skull with a scalpel, and the dura mater with sharp forceps. After injection or implant, the exposed brain surface was sealed with biocompatible adhesive (World Precision Instruments, KWIK-CAST). Birds were allowed to recover for at least one week before any data collection. For histology, brains were fixed in 4% paraformaldehyde solution for 2-3 days, then cut into 100-µm coronal slices with a vibratome and mounted on slides in polyvinyl alcohol (Sigma-Aldrich, 10981-100ML). Dark-field and fluorescent images were acquired using a fluorescence scope (Leica DM4000 B).

#### Tracer injection

To determine the stereotactic coordinates of MO/NAO and NLC, we first calculated a rough estimate based on the published budgerigar brain atlas (http://www.brauthlab.umd.edu/atlas.htm). We then injected 100 nL fluorescent cholera toxin subunit B (CTB, C34775, Invitrogen) tracers into the brain and measured the targeting accuracy with histology after 7-9 days. The coordinates were iteratively improved with 6 injections in 3 birds.

#### Microdialysis probe implant

We made custom microdialysis probes according to the method in Aronov et al 2008. The inflow polyimide tubing is approximately 2.25 mm long for MO/NAO implants and 4.25 mm long for the NLC implants. The cellulose membrane measures roughly 0.2 mm in width and 1 mm in length, with 0.7 mm exposed to deliver drugs. For MO/NAO, the probes were implanted bilaterally at 8.80 AP, 2.90 ML and 1.45 DV (unit: mm), with a 30° AP angle and 20° ML angle. For NLC, the probes were implanted bilaterally at 5.75 AP, 6.45 ML and 2.90 DV (unit: mm), with a 45° AP angle. To secure the implants, we inserted 4 insect pins into the skull and covered the base of probes with dental cement (C&B Metabond, Parkell, S380). The probes were flushed regularly with PBS to prevent clogging. After data collection, we infused fluorescein (10 mM, Krackeler Scientific, 45-46960-100G-F) into the probe to visualize implant location and drug diffusion.

#### Pharmacological inactivation

Compared to direct injection, a higher concentration of drug is needed with the microdialysis probe, since the solute needs to diffuse passively through the cellulose membrane (Aronov et al 2008). Following previous studies in the zebra finch (Aronov et al 2008), we diluted tetrodotoxin citrate (Abcam, ab120055-1MG) in PBS to 50 µM and muscimol (VWR, Q-1345.0050BA) in PBS to 1.5 mg/mL. One week after surgery, we started infusing PBS into probes to acclimate the bird to the procedure. Infusion was conducted roughly 1 hour before the cage light turned on. For each probe, we infused 1.0 mL solution at a rate of ∼0.2 mL/min. For TTX, we left the drug in the probe for 15 mins then washed it out with PBS. The bird was then placed back to the cage to rest before the light turned on and behavioral trials began. We used the same procedure for TTX and PBS, and conditions were alternated daily, so a direct comparison is meaningful. We had 2-3 infusion replicates for each bird under each condition. For experiments using muscimol, we diluted it to 1.5 mg/mL in PBS (Aronov et al 2008). Since muscimol degrades faster than TTX, to sample enough contact calls, we kept muscimol in the probes during data collection for about 6 hours, then washed it out with PBS.

#### Electrophysiological calibration

We calibrated the inactivation range of our reverse microdialysis probes using extracellular electrophysiological recordings. The primary goal of the calibration experiments was to show that we were able to inactivate MO/NAO while leaving distant structures intact. To demonstrate this convincingly, we needed a site with high baseline firing rates as a positive control to confirm that when a microdialysis probe was implanted nearby and infused with TTX, its neuronal activity was abolished. Our preliminary data suggested that MO/NAO has low baseline firing, which made it unsuitable to serve as the positive control. We thus conducted the calibration in adult zebra finches (>90 dph), taking advantage of the high baseline firing of area X and RA. We first mapped the locations of area X and RA using carbon fiber electrodes (Carbostar-1, Kation Scientific) and recorded the pre-implant activity with custom Matlab scripts. We then implanted a custom microdialysis probe in between, approximately ∼1 mm from area X and ∼4 mm from RA (Figure S1). We first infused PBS into the microdialysis probe with an extended Tygon tubing and 1 mL syringe and lowered the electrode to the pre-mapped area X site to record the pre-TTX activity. We then gently infused 1 mL TTX (50 µM) into the probe and recorded how the neuronal activity at the area X site changed with time. After 15 mins, we washed the TTX out with PBS, exactly as the procedures in our AFP inactivations experiments. Finally, we lowered the electrode to the pre-mapped RA site to record the neuronal activity.

### QUANTIFICATION AND STATISTICAL ANALYSIS

#### Contact call identification

We wrote custom Matlab scripts to identify contact calls from audio recordings. Noise at low and high frequency was first removed with a bandpass Butterworth filter (500-10,000 Hz). We then calculated the spectral flatness (Wiener energy) and set a threshold (0.65) to distinguish vocal signal from broadband noise. Segments that passed the threshold were considered candidates and exported for visual inspection. Contact call is characteristic in having a major modulated frequency band between 2-4 kHz. Cage noise, squawks, harmonic calls, and warble were removed from downstream analysis.

#### Acoustic feature analysis

We used the SAT Matlab package from Sound Analysis Pro 2011 to calculate acoustic features of identified calls (http://soundanalysispro.com/matlab-sat). Pitch modulation depth was defined as (max - min) / mean of pitch. Spectrograms were calculated using the Short-Time Fourier Transform (STFT) algorithm in Matlab (window size 512, nfft 512). Variability of acoustic features was calculated based on the method in Olveczky et al 2005, to facilitate direct comparison to results in zebra finches. Briefly, we randomly sampled 50 pairs of calls from each bird under each condition. Since subject F6 had two normal call types (Figure 5B), we only sampled call pairs that belonged to the same type. We then used the ‘Similarity batch’ function in Sound Analysis Pro 2011 to calculate pairwise similarity scores with the default parameters except for the ‘min interval’, which was set to 25 ms. The scores were calculated based on pitch, frequency modulation, amplitude modulation, Wiener entropy, and goodness of pitch. Following Olveczky et al 2005, values of ‘accuracy’ were used as the metric for similarity and further converted to variability scores (variability score = (100-similarity score)/50).

#### Variational auto-encoder (VAE)

We followed the method in Goffinet et al 2021 to train the VAE network for contact call analysis. Briefly, spectrograms of amplitude-normalized calls were centered in a 128*128 matrix and used as the inputs. We performed resampling so each individual had the same number of calls, then randomly split the dataset for training (80%) and testing (20%). The VAE architecture was adapted from Goffinet et al 2021 with dimension of latent features fixed to 32. The network was trained for 100 epochs at learning rate 0.001 and batch size 32 with the Adam optimizer implemented in Pytorch 1.12.1. We then calculated the latent features of each call by feeding it into the encoder network. Separate networks were trained for the behavioral (Figure 2) and inactivation experiments (Figure 3-5).

#### Uniform Manifold Approximation and Projection (UMAP)

We used the umap module in Python to visualize the VAE latent features of contact calls in low dimensional space (https://umap-learn.readthedocs.io/en/latest/index.html). The same parameters were used for all UMAP analyses (n_components=2, n_neighbors=20, min_dist=0.1, metric=’euclidean’, others=default). We have explored varying the n_neighbors and min_dist parameters, and obtained qualitatively similar results. We took special caution to use UMAP only as a visualization tool to provide intuitions. Quantifications were conducted using more rigorous methods, e.g. SVM. In the UMAP plots, because the number of calls varied across individuals, we set a max limit to achieve a more balanced representation of each individual. We used the *random* module in python to randomly select calls from each individual. No hand-picking was involved in this process. We also varied the random seed in the sampling procedure and obtained visually similar plots.

#### Support vector machine (SVM)

In past work, individual signatures in vocal signals were quantified with two major categories of approaches: 1) methods based on information theory ^60^; 2) classification models such as support vector machine (SVM) ^43,61^. Although information theoretic approaches allow comparison across different systems and quantification of features’ contribution to individuality, here our primary goal was to explicitly compare discriminability of the same system across conditions, e.g. train a model on the PBS dataset then check performance on the TTX dataset. We therefore adopted the classification approach to quantify individuality in contact calls. Specifically, we built SVM classifiers to decode individual identity from contact calls using the scikit-learn module in Python. We first conducted principal component analysis (PCA) on the VAE outputs to obtain the most informative features, then used them as inputs for SVM. We varied the dimension of principal inputs and calculated how the decoding accuracy changed accordingly. For each model, we sampled an equal number of calls from each individual to balance the dataset, then split it for training (80%) and testing (20%). Accuracy was calculated only on the testing dataset. The sampling procedure was repeated 50 times for each model. The mean accuracy and standard deviation were plotted in the figures.

#### Silhouette score calculation

We calculated the Silhouette score to quantify the separability of calls produced by different individuals. We first trained all calls produced on the PBS and TTX condition in a single VAE network, then reduced the dimension of latent features using PCA. We focused on the first 10 principal components, since together they could explain 97.8% of the total variance. We calculated the Silhouette score separately for the PBS and TTX dataset. We sampled 50 calls from each individual, then used the scikit-learn module in Python to calculate the Silhouette score on the first 10 principal components. The sampling was repeated 100 times to obtain a distribution of Silhouette scores. Shuffling of labels was also performed at each sampling to obtain a null distribution. To quantify the contribution of increased cross-rendition variability on the reduced discriminability, we normalized the standard deviation of individual clusters on each PCA axis, so the cluster sizes or spreads of each individual were the same for the PBS and TTX dataset. We then compared the Silhouette scores on the raw and standardized datasets.

### KEY RESOURCES TABLE

**Figure S1.**
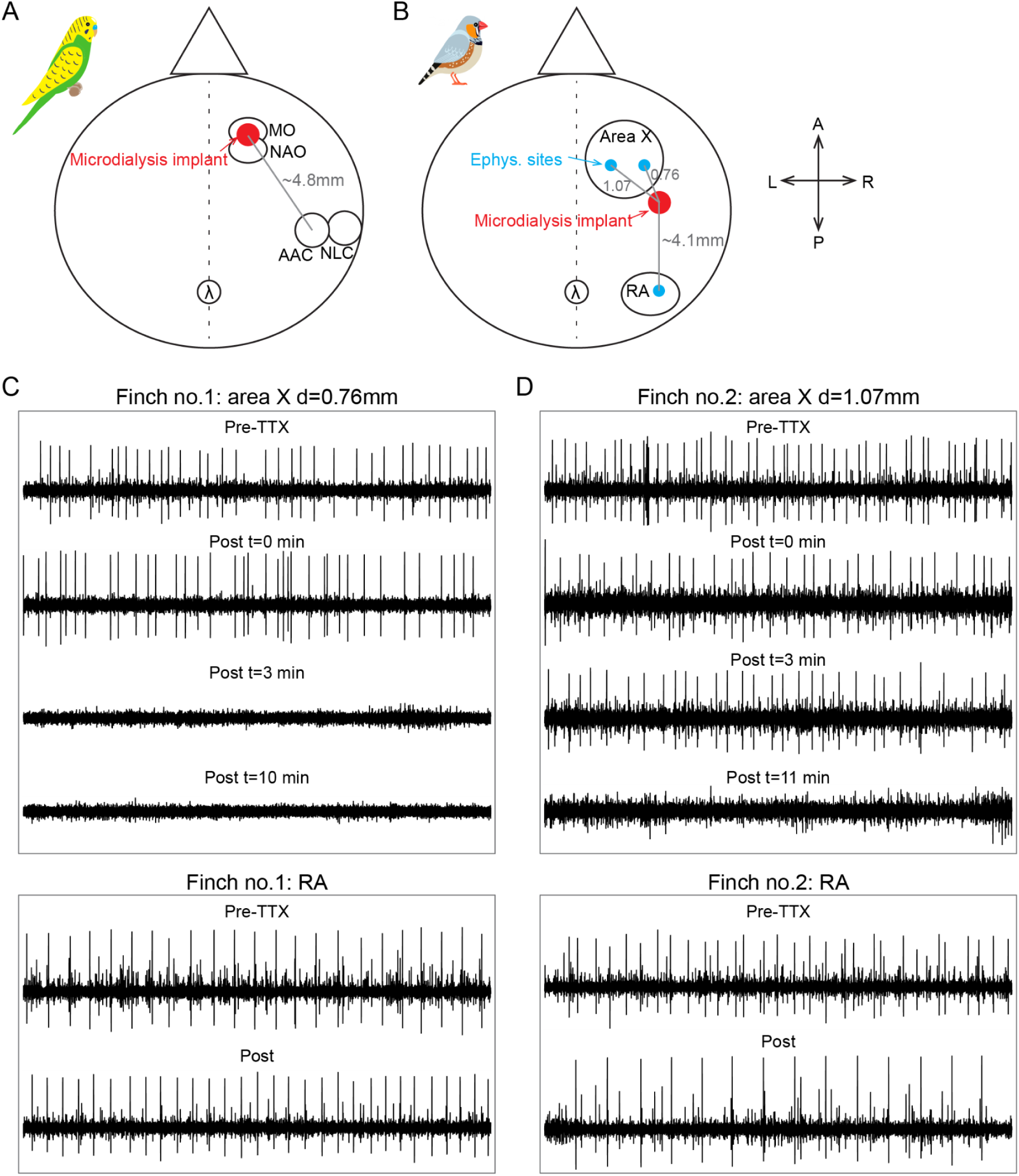
Calibration of inactivation range of reverse microdialysis probes. (A) Schematics showing the spatial relationship between the frontal microdialysis probes and the posterior NLC-AAC pathway in budgerigars, viewed from the top of the bird’s head. We estimated that the distance from the probe to the center of AAC was ∼4.8 mm. Note that we implanted probes bilaterally for the inactivation experiments, but only one side is shown here for clarity. (B) Because MO/NAO in budgerigars have low spontaneous activity under anesthesia, we carried out the calibration experiments in adult male zebra finches (>90 dph), taking advantage of the high spontaneous activity of area X and RA (Methods). We first mapped their locations using electrophysiology, then implanted a microdialysis probe between them, such that the probe is ∼1 mm from area X, while ∼4.1 mm from RA. We then infused TTX with the same concentration (50 µM) and volume (1 mL) as in the budgerigar AFP experiments, and monitored how the neuronal activity in area X and RA changed over time. (C) Results from one finch where the microdialysis probe was ∼0.76 mm from area X. We found that the spiking activity in area X disappeared rapidly within 3 mins (top), while activity in RA persisted even after 1 hour since TTX infusion (bottom). All example electrophysiological traces last 2.5 secs on the x-axis. Traces within the same box share the same y-axis. (D) Same as (C), but with a different finch where the microdialysis probe was ∼1.07 mm from area X. We observed similar results, except that area X activity lasted longer than (C), probably because it took the drug longer to travel to a more distant location. These calibration results demonstrated that with our inactivation method, we were able to cover the MO/NAO complex, while leaving the posterior NLC-AAC pathway intact.

**Figure S2.**
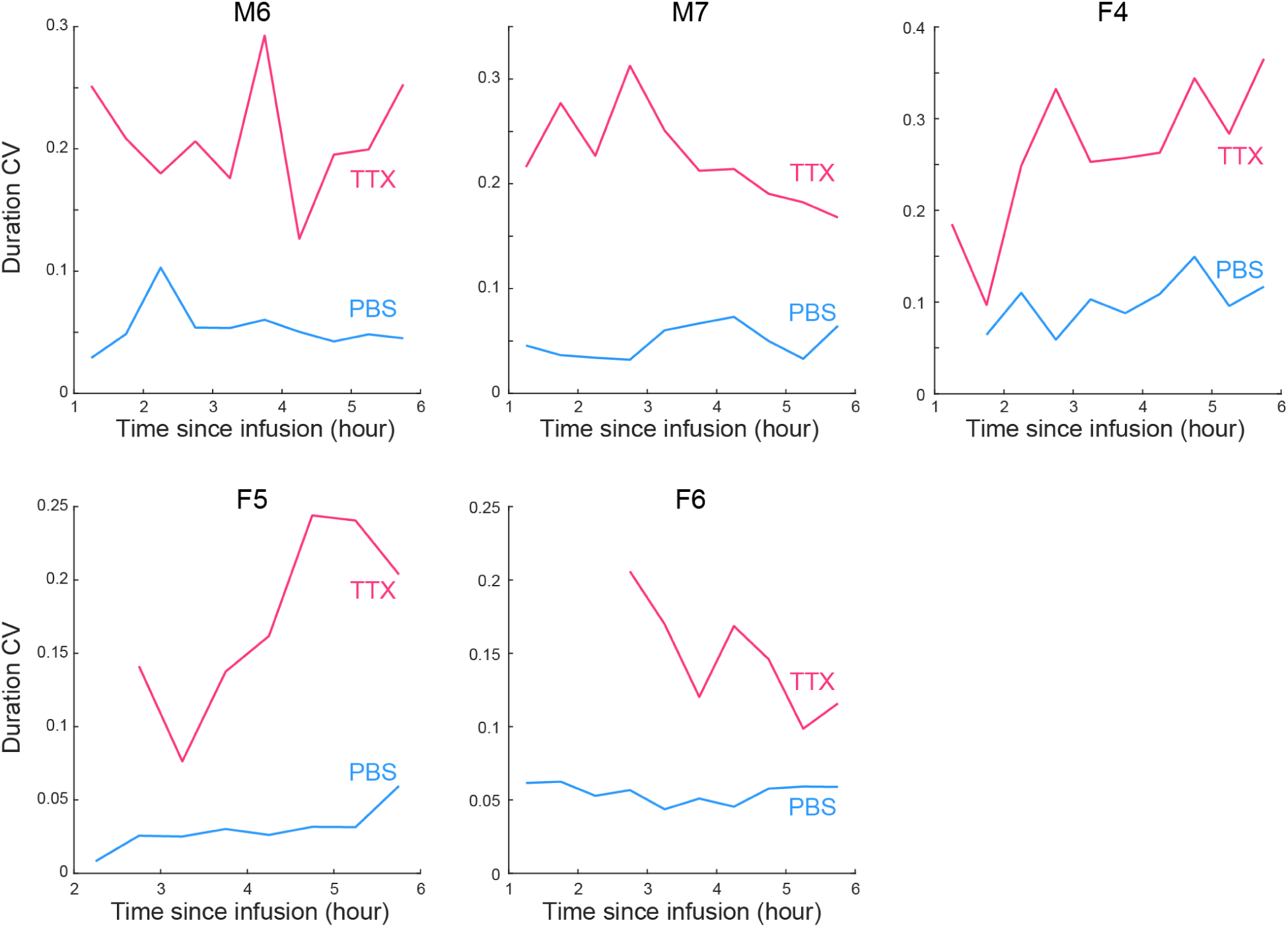
Cross-rendition variability is consistently higher for the TTX dataset over the entire experimental period. To check if the increased cross-rendition variability is an artifact of TTX losing effects over hours, we divided the experimental period since drug infusion into discrete time bins (30 mins), calculated variability of call duration within this short period, then examined how variability changes with time and if it differs between the PBS and TTX dataset. We found that the cross-rendition variability is higher at the TTX condition at all time bins for all birds, suggesting that the increased variability is not simply an artifact of TTX losing effects over time.

**Figure S3.**
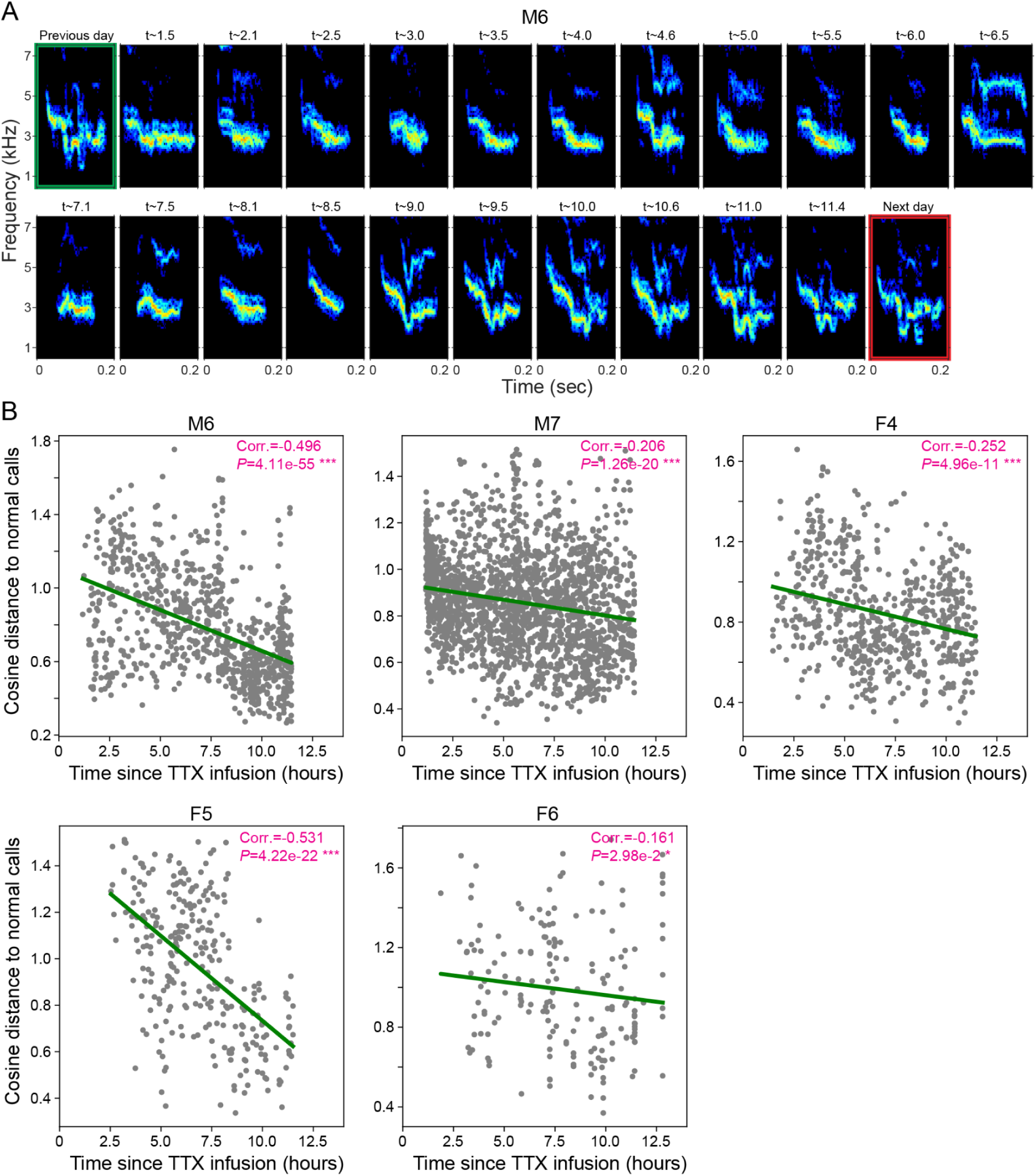
Contact calls gradually recover from AFP inactivation. (A) Spectrograms of contact calls from one example TTX infusion day. The green and red boxes show one call spectrogram from the same bird the day before and after TTX infusion, respectively. Numbers above other spectrograms indicate the time since TTX infusion in the unit of hour. Consistent with previous studies, the contact calls began to recover gradually after 9-10 hours of TTX infusion. (B) Quantification of the recovery process. We trained a single VAE network on contact calls produced under the PBS and TTX condition to obtain latent representations (dim=32), then calculated the dissimilarity (cosine distance) of each TTX call to the center of all PBS calls. Each plot is one bird, each gray dot is one call. We found a significantly negative correlation between dissimilarity and time in all birds, i.e. the calls became more similar to normal PBS calls over time. Values in pink indicate the Spearman correlation coefficients and p-values. Green lines indicate the prediction from a linear regression model.

**Figure S4.**
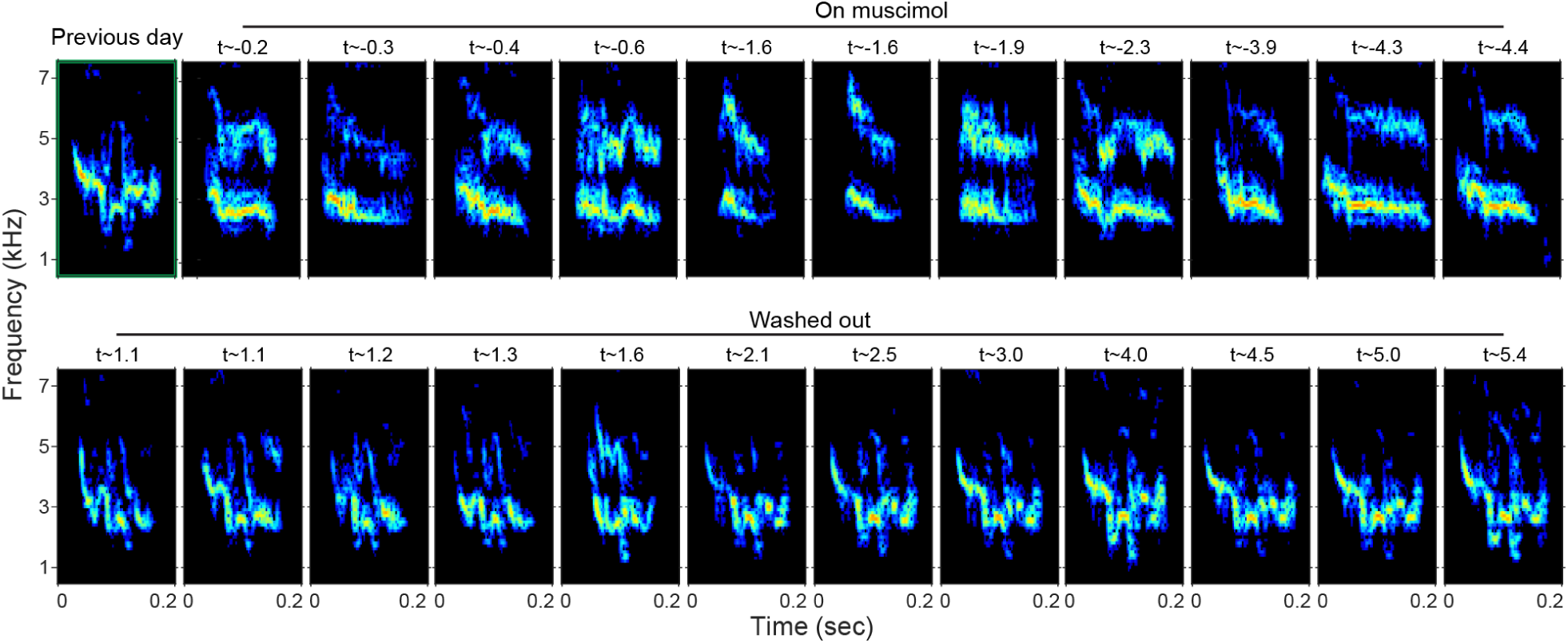
Inactivation by muscimol had a similar effect on contact call acoustic structure. Spectrograms of contact calls from one example muscimol infusion trial. The green box shows one call spectrogram from the same bird the day before muscimol infusion. Numbers above spectrograms indicate the time in hours since muscimol was infused in (top row) or washed out (bottom row) with saline.

**Figure S5.**
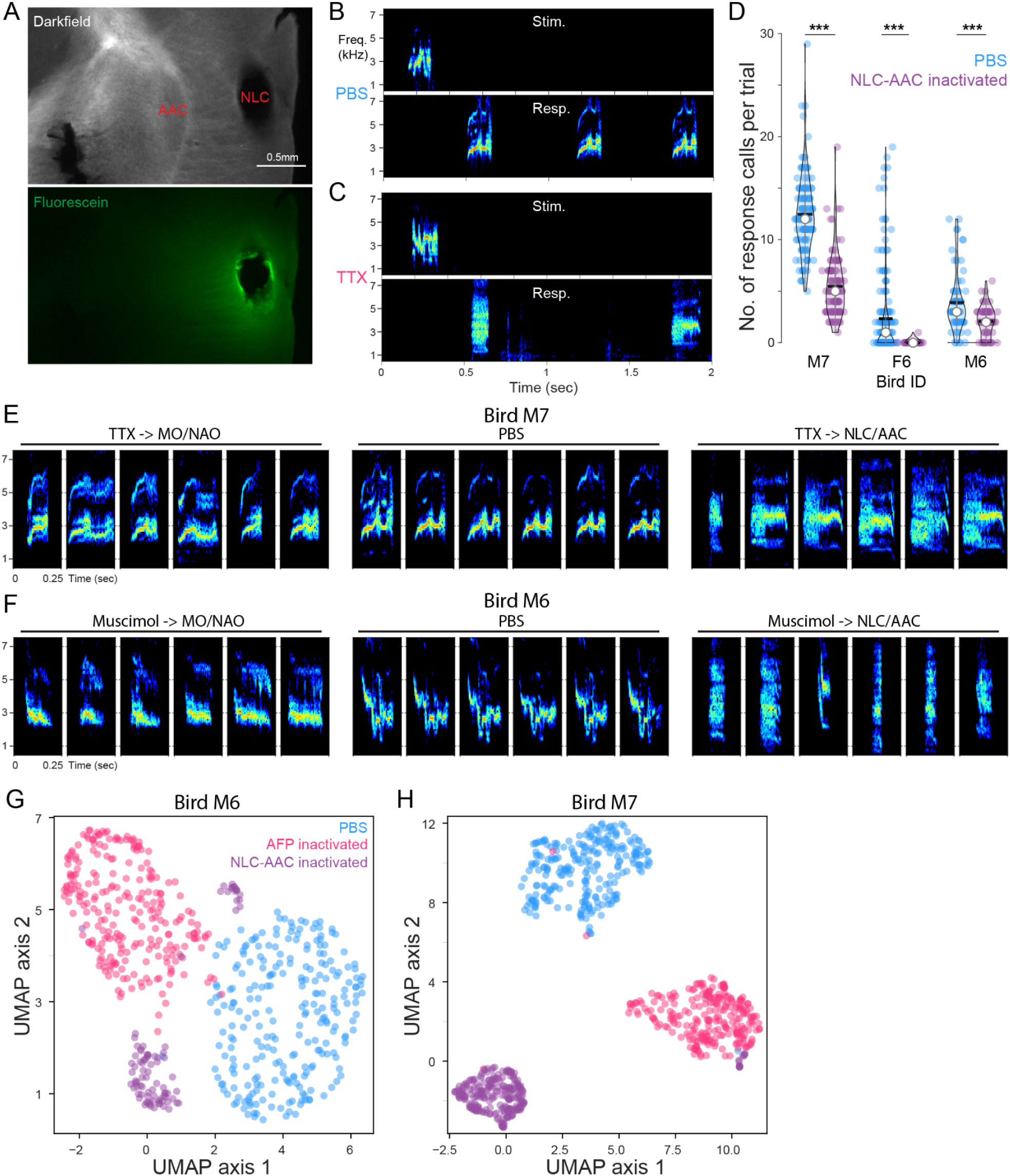
Effects of NLC-AAC inactivation on the production of contact calls. (A) Visualization of drug diffusion in the NLC inactivation experiments. We targeted bilateral NLC, but because AAC immediately neighbors NLC, the drug might also diffuse into AAC. (B) Spectrograms showing one example trial under the PBS condition. Top: stimulus; bottom: responses. (C) Same as (B), but for the TTX condition. When birds responded to the stimulus, they used vocalizations that contained broad-spectrum noise bands. (D) Response rates decreased significantly in all three birds. Each colored dot is one trial. The violins indicate the shape of distribution. The black line and white dot indicate the mean and median. P-value: ***<0.001, with Mann-Whitney U tests. (E) Side-by-side comparison of the effects of AFP inactivation (left) and NLC-AAC inactivation (right) in the same bird. Spectrograms of normal contact calls are shown in the middle. When AFP was inactivated, the frequency modulation was degraded and call duration became more variable, but the major frequency band was still centered around 3 kHz. However, when NLC-AAC was inactivated, the birds responded with calls that had broad-spectrum noise bands, which were more similar to the alarm calls. (F) Same as (E), but for a different bird with muscimol inactivation. (G-H) Embedding of contact calls produced under the PBS (blue), AFP-inactivation (pink), and NLC-AAC inactivation (purple) condition in the UMAP space of two different birds. Note that the cluster sizes in UMAP space don’t correspond directly to acoustic variability.

**Figure S6.**
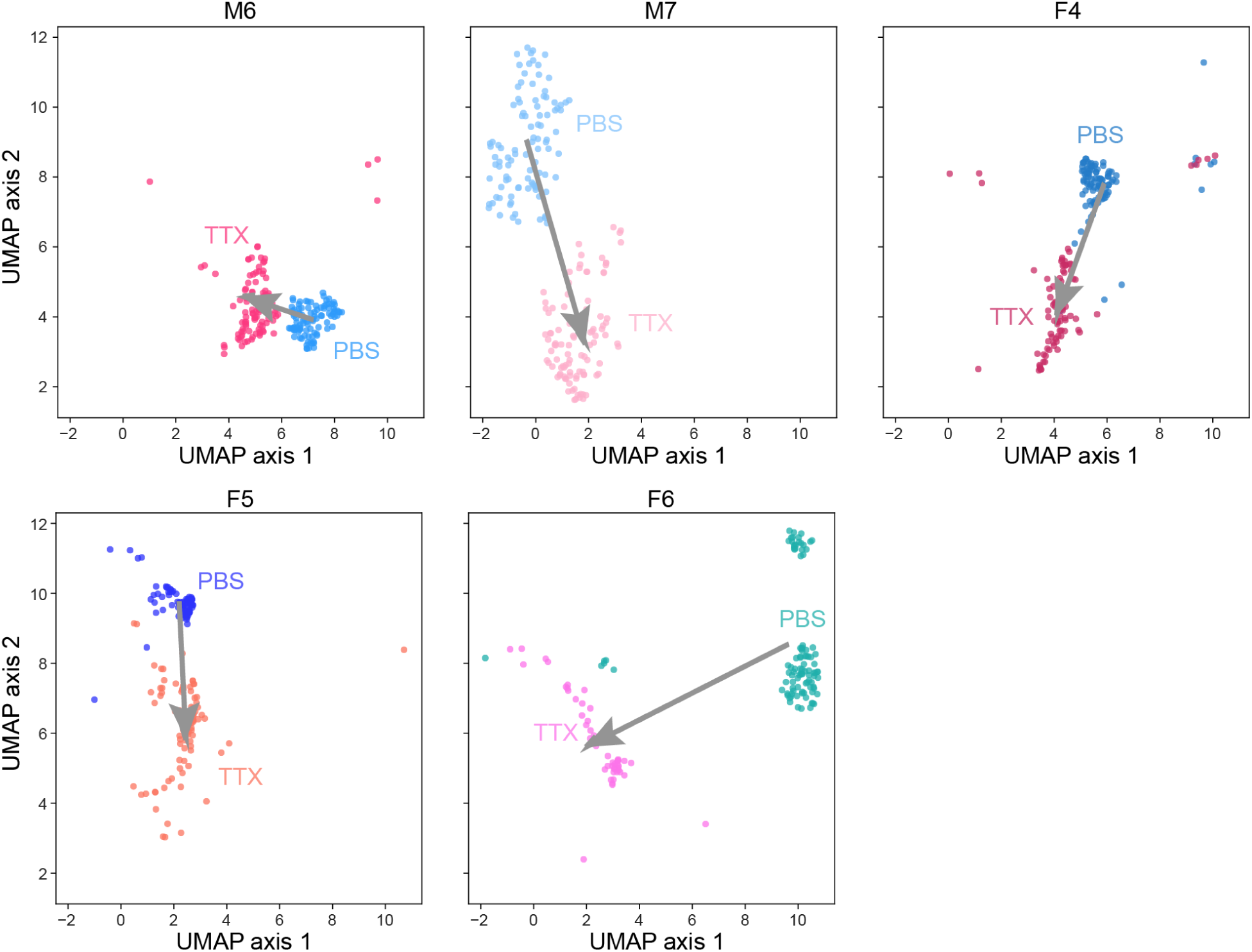
UMAP embedding of contact calls produced under the normal and AFP inactivation condition. Replot of the same data in Figure 5B by splitting each bird into one figure panel. Each colored dot is one sampled contact call. Arrows point from the PBS cluster centers to the TTX cluster centers.

